# On the Development of Sesamoid Bones

**DOI:** 10.1101/316901

**Authors:** Shai Eyal, Sarah Rubin, Sharon Krief, Lihi Levin, Elazar Zelzer

**Affiliations:** Weizmann Institute of Science, Department of Molecular Genetics, PO Box 26, Rehovot 76100, Israel

**Keywords:** Sesamoid bone, Patella, Fabella, Digits, Sox9, Scleraxis, Tgfβ, Bmp2, Bmp4, Joint, Mouse

## Abstract

Sesamoid bones are a special group of small auxiliary bones that form in proximity to joints and contribute to their stability and function. Sesamoid bones display high degree of variability in size, location, penetrance and anatomical connection to the main skeleton across vertebrate species. Therefore, providing a comprehensive developmental model or classification system for sesamoid bones is challenging. Here, we examine the developmental mechanisms of three anatomically different sesamoid bones, namely patella, lateral fabella and digit sesamoids. Through a comprehensive comparative analysis at the cellular, molecular and mechanical levels, we demonstrate that all three types of sesamoid bones originated from *Sox9^+^/Scx^+^* progenitors under the regulation of TGFβ and independent of mechanical stimuli from muscles. We show that BMP4 was necessary specifically for differentiation of patella but not of lateral fabella or digit sesamoids, whereas BMP2 regulated the growth of all examined sesamoids. Next, we show that whereas patella and digit sesamoids initially formed in juxtaposition to long bones, the lateral fabella formed independently at a distance. Finally, we provide evidence suggesting that while patella detached from the femur by formation of a synovial joint, digit sesamoids detached from the phalanx by a fibrocartilage joint. Collectively, these findings highlight both common and divergent molecular and mechanical features of sesamoid bone development, thereby advancing our understanding of their evolutionary plasticity.

## INTRODUCTION

Sesamoid bones constitute a special group of bones that are integral to the skeletons of many vertebrate species (Abdala et al., 2017; Bizarro, 1921; Corina Vera et al., 2015; Jerez et al., 2009; Maisano, 2002; Ponssa et al., 2010; Sarin et al., 1999). Estimated to have evolved 200 million years ago (Carter, 1998), sesamoid bones are usually small and flat with morphological resemblance to the sesame seed, hence their name (Vickaryous and Olson, 2007). In addition, sesamoid bones are typically embedded within tendons, notably in proximity to joints. However, despite these common characteristics, sesamoids vary tremendously in aspects such as location and penetrance both within and across species, and the type of joints they are associated with. This high degree of variability imposes a challenge on providing a comprehensive model for sesamoid bone development.

The patella, also known as the kneecap, is the largest sesamoid bone in mammalians and is part of the patellofemoral joint, one of two joints composing the knee (Samuels et al., 2017). The patella is positioned within the patellar groove located at the anterodistal part of the femur and is separated from it by the synovial patellofemoral joint. The patellar groove enables the patella a controlled slide across during flexion and extension of the lower leg segment, whereas the synovial joint permits this high degree of flexibility while maintaining the integrity of the articular cartilage of the long bones flanking the synovial capsule (Schindler and Scott, 2011). The patella has a crucial effect on the stability and mechanics of the knee. It absorbs disruptive forces applied to the quadriceps tendon, within which it is embedded, and enhances the moment arm and resulting extension force of the quadriceps muscle (Bizarro, 1921; Lennox et al., 1994; Mottershead, 1988; Sarin et al., 1999; Sutton et al., 1976).

The smaller fabella is located opposite the patella at the back of the mammalian knee. Although it is similarly embedded within the lateral head tendon of the gastrocnemius muscle in both human and mouse, the attachment site of this tendon varies between the two species. Thus, in humans the fabella is located behind the lateral condyle of the femur, whereas in mice it is found on its lateral side (Jin et al., 2017; Phukubye and Oyedele, 2011; Sarin et al., 1999). The fabella lacks articular cartilage and is additionally connected by the fabellofibular ligament to the fibular tip and to the proximolateral side of the meniscus by two separate bundles of fibers (Jin et al., 2017). These anatomical differences imply that unlike patella, the function of the fabella is not to facilitate and enhance movement but rather restrict it by anchoring and stabilizing the knee, thus preventing possible external rotation of the tibia or hyperextension of the knee (Hauser et al., 2015).

In mice, digit sesamoids can be found in pairs at any metacarpo/metatarsophalangeal and proximal interphalangeal joints of any digit, where they are dorsally embedded within the flexor digitorum tendon and anteriorly share articulation with the metacarpal/metatarsal bones. In contrast to the patella, they are unable to freely glide over these articulated joints, as they are physically connected to the proximal tip of the proximal phalanx by a fibrocartilaginous joint (Doherty et al., 2010). Thus, digit sesamoids serve as a guide for the long flexor tendons in directing and controlling the course of the strings during flexion of the digits (Wirtschafter and Tsujimura, 1961).

To date, the prevailing model for sesamoid development suggests that these bones are induced within the tendons that wrap around joints by mechanical forces generated by embryonic movement (Hall, 2005; Parsons, 1904). However, we recently reported that patella forms by a different developmental mechanism (Eyal et al., 2015). Briefly, we demonstrated that patella originates from a distinct pool of *Sox9* and scleraxis (*Scx*) double-positive precursors in juxtaposition to the femur, which are later separated from it by the application of a joint formation program. TGFβ and BMP4 signaling pathways were shown to be required for patella precursor specification and differentiation, respectively. Finally, we found that mechanical load is not necessary for patella formation, but is needed for its separation from the femur by regulating the formation of the patellofemoral joint. Although our findings pertain specifically to patella formation, they raise the question of whether our model for patella development could be applied to other sesamoid bones.

In this study, we test our model of patella development on the development of the fabella and digit sesamoids. We show that all examined sesamoid bones originated from *Sox9*^+^*/Scx*^+^ progenitors that were induced independently of mechanical stimuli. We further show that these progenitors require TGFβ signaling for their specification and BMP2 signaling for their growth, whereas BMP4 signaling is required for patella differentiation only. We provide evidence suggesting that sesamoids may develop away from the skeleton, or separate from the bone shaft through mechanical load-dependent formation of a synovial joint, or through mechanical load-independent formation of a fibrocartilaginous joint.

## RESULTS

### *Sox9*^+^*/Scx^+^* chondroprogenitors give rise to sesamoid bones

Previously, we described a mechanism for patella development (Eyal et al., 2015). To address the question of the generality of this mechanism, we analyzed comparatively patella development with that of two other subgroups of sesamoid bones, namely the lateral fabella and the metacarpophalangeal sesamoids. Like the patella, these sesamoids display full penetrance in mice. We first compared the anatomy of the three sesamoid bones by both micro-CT scans and histological sections (Fig. 1A-E). Our results showed that by E18.5, all sesamoid bones have formed and were embedded within tendons. The patella, which was separated from the femur by the patellofemoral synovial joint on its dorsal side, was ventrally embedded within the quadriceps tendon (Fig. 1A, Ci and Cii). The lateral fabella (LF) was embedded within the tendon of the lateral head of the gastrocnemius muscle, which attaches the dorsolateral condyle of the femur, and connected at its base to the lateral meniscus by the fabellofibular ligament (Fig. 1A, Di and Dii). The third metacarpophalangeal sesamoid bone (III-MCPS) was separated from the metacarpal (MC) bone by synovial space on its ventral aspect and embedded within the flexor digitorum tendon (FDT) on its proximodorsal aspect (Fig. 1A, Ei and Eii). Micro-CT imaging showed that the III-MCPS was distinct from the proximal phalanx (PP); yet, histologically we observed an unfamiliar tissue connecting between the III-MCPS and the posterior tip of the proximal phalanx (Fig. 1Eii; enlarged rectangle), most likely the primordium of the fibrocartilage tissue that connects the MCPS to the PP in adult mice (Doherty et al., 2010). Patella precursors, which co-express *Sox9* and *Scx*, are added secondarily to the *Sox9^+^/Col2a1^+^* chondrocytes of the femur anlage (Eyal et al., 2015). To examine whether the LF and III-MCPS originated from similar progenitors, we first sought to find the developmental stage at which these sesamoids start to form. For that, we harvested and sectioned embryonic WT knees or carpal digits at sequential developmental stages from E13.5 to E17.5 and immunostained them using antibodies for SOX9, a marker for chondroprogenitors, and COL2A1 as a marker for chondrocytes. At E14.5, *Sox9^+^/Col2a1^-^* cells marked the undifferentiated LF precursors, whereas *Sox9^+^/Col2a1^+^* differentiated chondrocytes marked the femur and tibia (Fig. 1Diii). Notably, whereas the *Sox9*^+^*/Col2a1*^-^ patella precursors developed in juxtaposition to the femur, LF precursors were not connected to either femur or tibia (Fig. 1Ciii and Diii).

**Figure 1.**
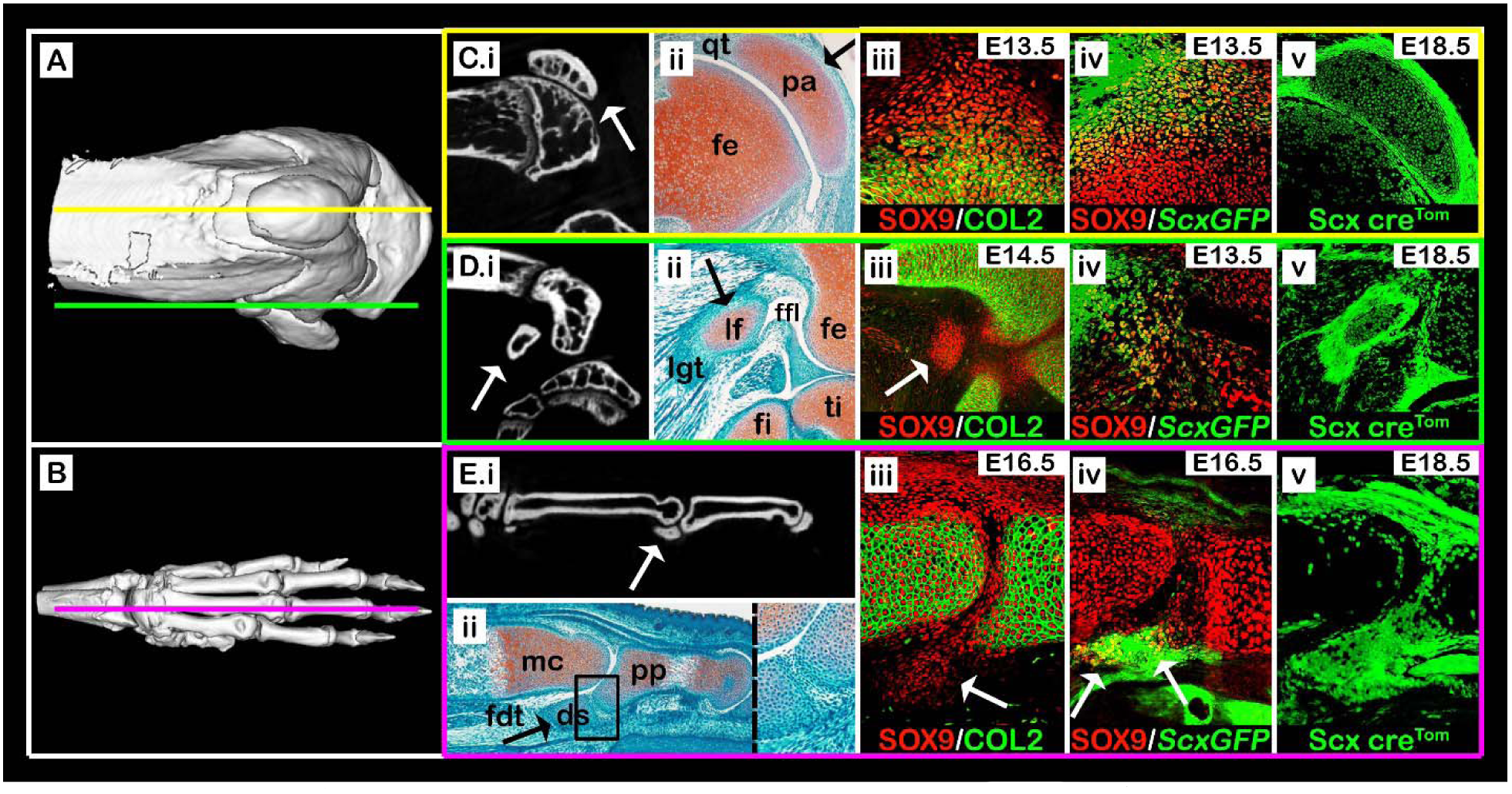
Sesamoid bones arise from *Sox9^+^/Scx^+^* progenitors. **(A,B)** Micro-CT scans of an adult knee (A) and forelimb autopod (B). Yellow line (A) indicates sagittal sections through the patella, as shown in panels C(i-v), green line (A) indicates sections through the lateral fabella, as in panels D(i-v), and purple line (B) indicates sections through the digit sesamoid, as in panels E(i-v). **(C.i-E.i)** Selected sagittal planes from micro-CT scans of both knee and autopod; white arrows indicate sesamoid bones. **(C.ii-E.ii)** Sagittal sections through limbs of E17.5 WT embryos stained by Safranin-O. Black arrows indicate sesamoid bones. (C.ii): The patella (pa) is embedded within the quadriceps tendon (qt). (D.ii): The lateral fabella (lf) is embedded within the lateral gastrocnemius tendon (lgt). (E.ii): The digit sesamoid (ds) is embedded within the flexor digitorum tendon (fdt) and connected to the proximal phalanx (pp) by a fibrocartilage tissue (enlarged rectangle). **(C.iii-E.iii)** Sagittal sections through limbs of WT embryos immunostained against SOX9 and COL2A1 show that sesamoid bones are formed at different developmental stages after the differentiation of the long bone cartilaginous anlagen (white arrows). Notice the specification of patella and digit sesamoid precursors in continuum with the bone shaft anlagen. **(C.iv-E.iv)** Sagittal sections through limbs of *ScxGFP* embryos stained against SOX9 show that all three sesamoid bones are formed by *Sox9*^+^*/Scx*^+^ progenitors (white arrows). **(C.v-E.v)** Sagittal sections through E18.5 limbs from *Scx-Cre^tdTomato^* embryos indicate that all three sesamoid bones were formed by precursor cells that descended from *Scx*-expressing cells. Abbreviations: fe, femur; ti, tibia; fi, fibula; mc, metacarpal bone.

The *Sox9^+^/Col2a1^-^* III-MCPS precursors were first observed at E16.5, subsequent to the differentiation of both MC and PP anlagen. Interestingly, at that stage, these precursors extended continuously from the proximodorsal aspect of the PP (Fig. 1Eiii). This implied that later in development, they would have to detach from the PP, similarly to the patella.

Having established the initiation stage of these sesamoids, we next examined their cellular origin. For that, sectioned knees and carpal digits of transgenic *ScxGFP* embryos were immunostained for SOX9 to highlight chondrocytes and, specifically, *Sox9*^+^*/Scx^+^* chondroprogenitors. We observed large populations of *Sox9^+^/Scx^+^* chondroprogenitors at the presumptive sites of patella, LF and III-MCPS formation (Fig. 1Civ, Div and Eiv). To further confirm that both LF and III-MCPS cells were descendants of *Scx^+^* chondroprogenitors, we performed a lineage tracing experiment using the *Scx*-*Cre* driver mouse crossed with tdTomato reporter mouse. As expected, we observed in sagittal sections of E18.5 knees and forelimb digits that patella, LF and III-MCPS cells were of *Scx^+^* lineage (Fig. 1Cv, Dv and Ev, respectively). Collectively, these results confirmed that both LF and III-MCPS precursors formed in accordance to the patella model, namely from distinct *Sox9*^+^*/Scx*^+^ chondroprogenitors that formed following differentiation of the long bone primordium cells.

### Initiation of sesamoid bones is independent of mechanical load

Sesamoid bone development has been assumed to be induced by mechanical signals produced by embryonic movement (Parsons, 1904; Parsons, 1908). Yet, we have reported that patella induction was mechanical load-independent (Eyal et al., 2015). We therefore reexamined the paradigm of mechanical load-dependent induction of sesamoid bones by studying the formation of LF and III-MCPS in paralyzed *mdg* mutant embryos (Pai, 1965). Examination of histological sections through knees and carpal bones of E17.5 mutants showed, as was previously described, that patella was developing (Fig. 2A, A’). Similarly, we observed that both LF and III-MCPS were formed in the mutant as well (Fig. 2B-C’). These results provide conclusive support for our model by indicating that the induction of sesamoid bone formation is independent from muscle-induced mechanical load.

**Figure 2.**
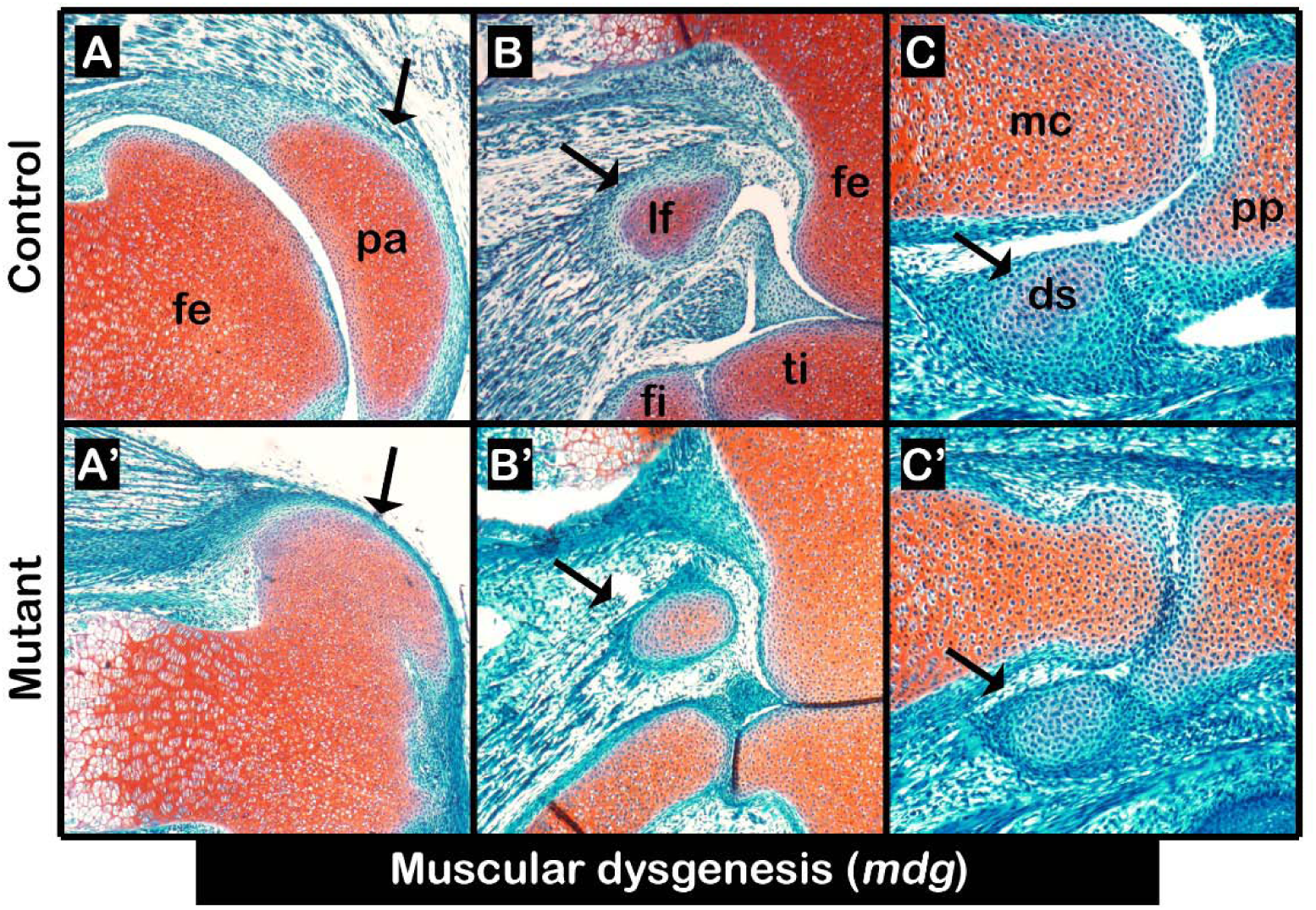
Initiation of sesamoid bone formation is independent of mechanical stimuli. **(A-C’)** Sagittal sections through E17.5 limbs from *mdg* control (A-C) and mutant (A’-C’) embryos stained by Safranin-O show that mechanical load is not needed for sesamoid bone initiation or growth, nor for fibrocartilage joint formation between digit sesamoid and proximal phalanx; yet, it was needed for patellofemoral joint formation. Black arrows indicate sesamoid bones. Abbreviations: fe, femur; pa, patella; lf, lateral fabella; ti, tibia; fi, fibula; mc, metacarpal bone; pp, proximal phalanx; ds, digit sesamoid.

### Initiation of lateral fabella and third metacarpophalangeal sesamoid bones is regulated by TGFβ but not BMP signaling

Another feature of patella development is the regulation of its *Sox9*^+^*/Scx*^+^ precursors by both TGFβ and BMP4 signaling pathways (Eyal et al., 2015). Having identified a similar cellular origin, we proceeded to examine the involvement of these pathways in LF and III-MCPS development. Additionally, we examined the involvement of *Bmp2,* a paralog of *Bmp4* (Pignatti et al., 2014). For that, *Tgfbr2, Bmp4* or *Bmp2* were specifically knocked out (cKO) from early limb mesenchyme using *Prx1*-*Cre* as a deleter mouse (*Prx1-TgfβRII^floxed^; Prx1-Bmp4^floxed^; Prx1-Bmp2^floxed^*) (Chytil et al., 2002; Liu et al., 2004; Logan et al., 2002; Ma and Martin, 2005; Selever et al., 2004). Then, sections through the knees and digit bones of E17.5 mutant and control embryos were stained by Safranin-O (Fig. 3A-F”). Our examination revealed that similarly to patella, both LF and III-MCPS were completely absent in *Prx1-TgfβRII^floxed^* mutants (Fig. 3A-B”). Surprisingly, we found that the loss of *Bmp4* in limb mesenchyme resulted only in patella aplasia, but had no effect on the formation of LF and III-MCPS (Fig. 3C-D”). Finally, examination of the *Prx1-Bmp2^floxed^* mutants revealed that initiation of all sesamoids was independent of BMP2 signaling; however, all sesamoids displayed growth retardation (Fig. 3E-F”). Collectively, these results suggest that sesamoid bones are regulated globally by TGFβ signaling, which drives their initiation, and by BMP2 signaling controlling growth. By contrast, BMP4 signaling was necessary specifically for patella formation but not for that of other sesamoids.

**Figure 3.**
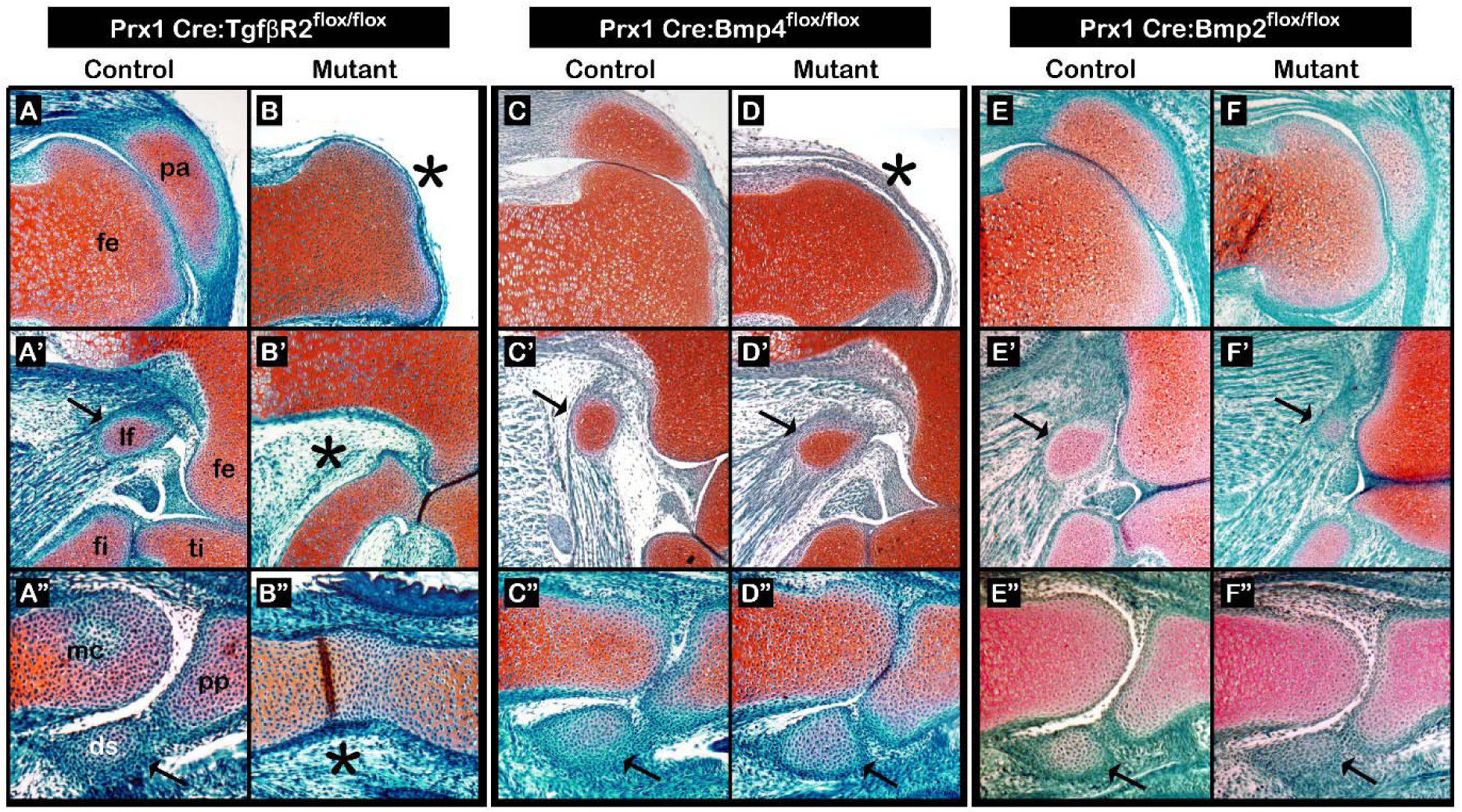
TGFβ signaling regulates initiation of all sesamoid bones, whereas BMP signaling has varying regulatory roles. Sagittal sections through limbs from E17.5 *Prx1-TgfβRII^floxed^* (A-B”), *Prx1-Bmp4^floxed^* (C-D”), and *Prx1-Bmp2^floxed^* (E-F”) mutant and control embryos stained by Safranin-O. **(A-B”)** Limb-specific ablation of TGFβ signaling resulted in sesamoid bone aplasia (B-B”). Black asterisks indicate missing sesamoid bones. **(C-D”)** Limb-specific ablation of *Bmp4* resulted in patellar aplasia, whereas both LF and III-MCPS appear unaffected. **(E-F”)** Limb-specific ablation of *Bmp2* did not affect sesamoid bone initiation, yet resulted in sesamoid bone hypoplasia. Abbreviations: fe, femur; pa, patella; lf, lateral fabella; ti, tibia; fi, fibula; mc, metacarpal bone; pp, proximal phalanx; ds, digit sesamoid.

### The metacarpophalangeal sesamoids are separated from the phalanx tip by formation of a fibrocartilaginous joint

Although digit sesamoids are distinct from the proximal phalanx in neonatal mice, at E16.5 the III-MCPS precursors appeared to be continuous with the cartilage anlage of the PP (Fig. 1Eiii). By E18.5, these precursors were detached from the PP while remaining connected to it by a fibrocartilage-like tissue (Fig. 1Eii). We hypothesized that the detachment process was mediated by formation of a fibrocartilaginous joint. To examine this possibility, we performed double-fluorescent in situ hybridization using probes for cartilage marker *Sox9,* tendon cell markers *Scx* and *Tnmd*, and joint-forming cell markers *Gdf5* and *Tppp3*. As expected, *Scx* and *Tnmd* expressions were observed within the tendon enveloping the patella on its ventral aspect, whereas *Gdf5* and *Tppp3* were expressed within the forming patellofemoral joint (Fig. 4A-D). Within the tissue connecting the MCPS and the proximal phalanx *Scx* expression was observed (Fig. 4A’), whereas expression of *Tnmd*, *Gdf5* or *Tppp3* could not be detected (Fig. 4B’-D). Because *Tnmd* was shown to be expressed in differentiating tenocytes (Shukunami et al., 2006), the expression of *Scx* but not *Tnmd* in this connective tissue suggests that it would not proceed to differentiate into a tendon or ligament. Instead, it could suggest the development of a fibrocartilaginous joint.

**Figure 4.**
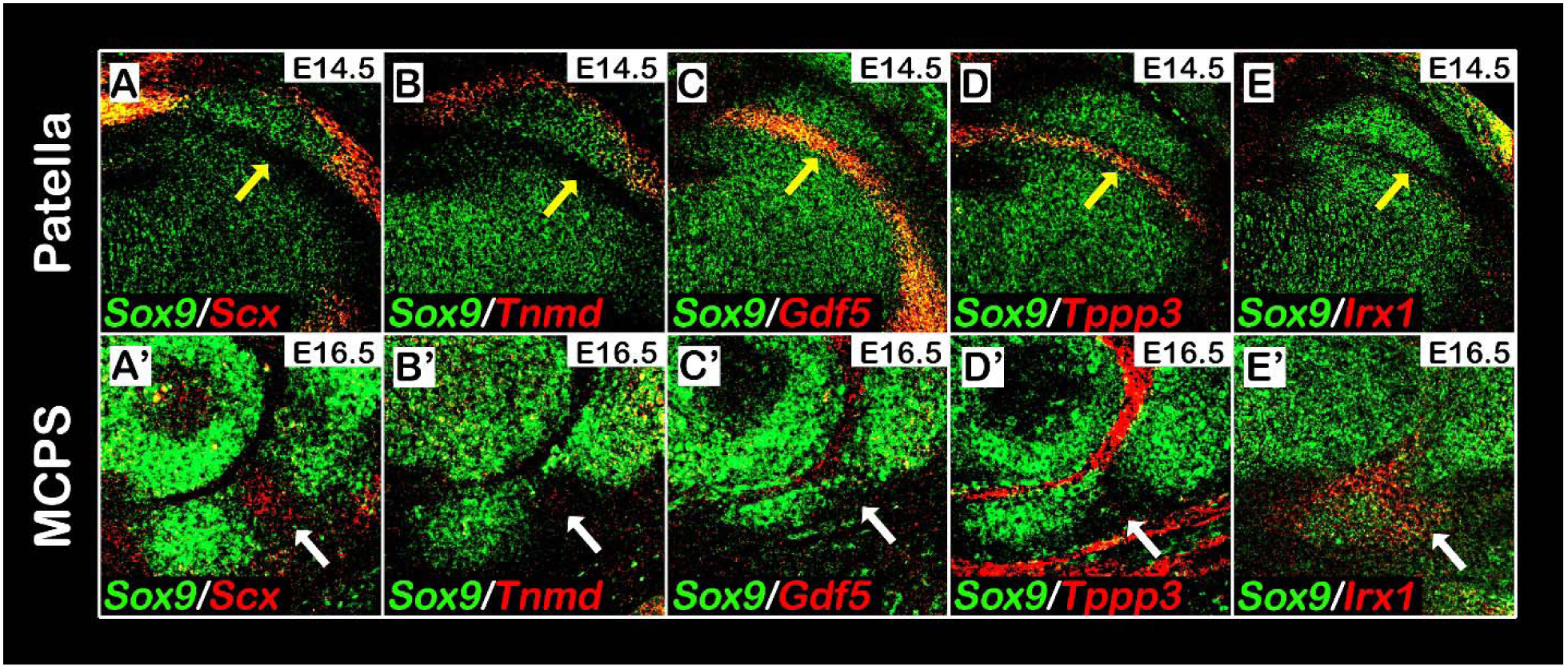
*Irx1* expression indicates the formation of a fibrocartilaginous joint separating the metacarpophalangeal sesamoid from the proximal phalanx. **(A-E’)** Sagittal sections through E14.5 knees (A-E) or E16.5 digits (A’-E’) from WT embryos analyzed by double fluorescence in situ hybridization using digoxigenin-and fluorescein-labeled antisense RNA probes for *Sox9* and *Scx, Tnmd, Gdf5, Tppp3* or *Irx1*. The patella is ventrally embedded within the quadriceps tendon, marked by *Scx* and *Tnmd* expression (A,B), and separated dorsally by the patellofemoral synovial joint, marked by *Gdf5* and *Tppp3* expression (C,D). In contrast, III-MCPS is neither connected to the proximal phalanx by a tendon nor separated from it by a synovial joint from, as indicated by the absence of *Tnmd* (B’), *Gdf5* (C’) and *Tppp3* (D’) expression at the connecting fibrocartilage tissue (marked by white arrows). (E-E’): Analysis of *Irx1* expression indicates that whereas *Irx1* is highly expressed in the fibrocartilage tissue (E’), it is completely absent from the patellofemoral synovial joint region (E).

**Figure 5.**
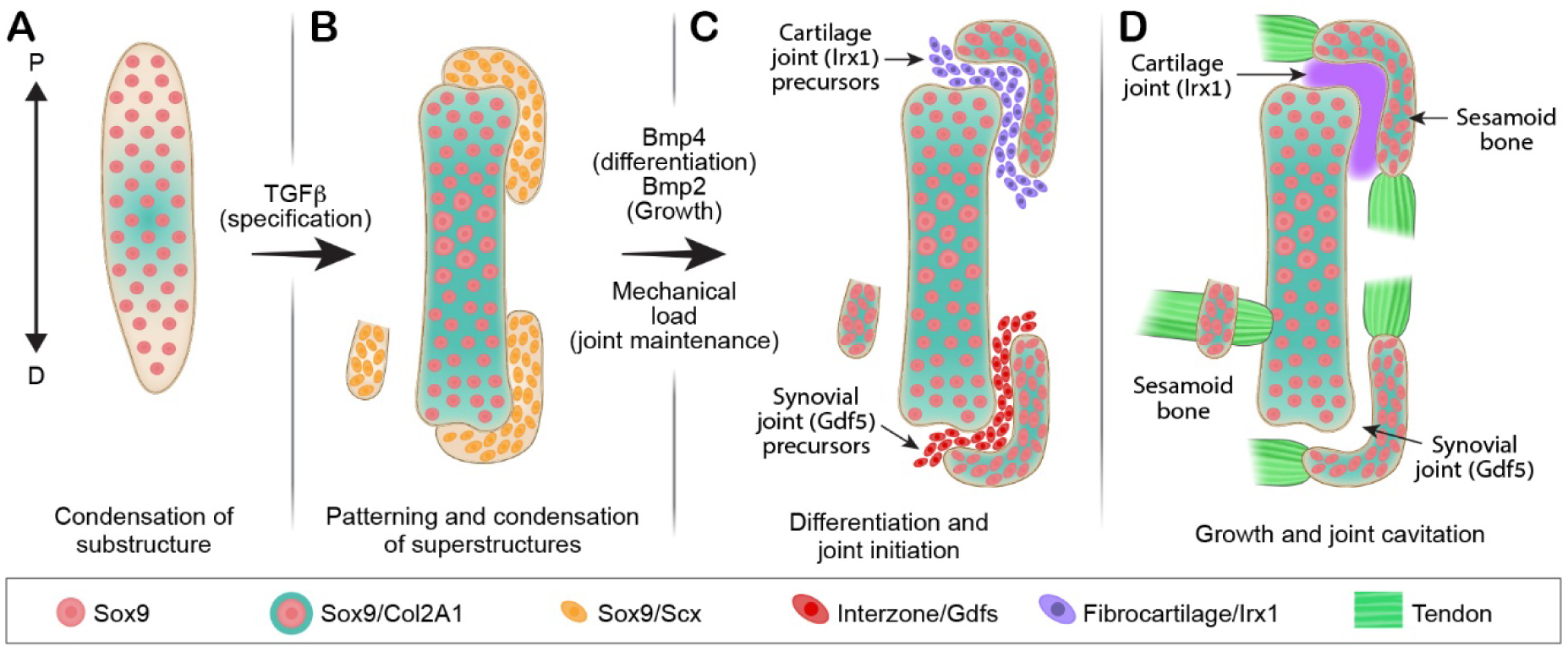
Molecular and mechanical signals facilitate development and anatomical variation of sesamoid bones. **(A,B)** We propose that secondary pools of *Sox9* and *Scx* co-expressing progenitor cells are specified in proximity to the primary *Sox9^+^* cartilaginous anlagen. These cells are regulated by TGFβ and BMP4 that are required for their specification and differentiation, whereas BMP2 signaling is required for sesamoid bone growth. **(C)** After differentiation, some of these modules become bone eminences that are integral to the skeletal substructure and serve as attachment sites for tendons, whereas others develop away from it and form a sesamoid bone. However, under different environmental conditions and pressures, attached modules can be separated from the cartilage anlage by formation of a joint between them. This leaves the module superficially embedded within the tendon, thus creating a sesamoid bone **(D)**.

Recently, Iroquois genes *irx5a* and *irx7* where shown to mark the forming cartilaginous hyoid joint in the zebrafish (Askary et al., 2015), whereas *Irx1* was reported to be highly expressed within the digit joints in mice (McDonald et al., 2010). We therefore examined the involvement of Irx genes in sesamoid bone development. For that, we performed double-fluorescent in situ hybridization using complementary RNA probes against *Sox9* and *Irx1.* As can be observed in Fig. 4E’, *Irx1* was specifically expressed within the connecting tissue between the MCPS and the PP, but was excluded from other joint compartments in the digits and from the patellofemoral synovial joint (Fig. 4E, E’). The combined expression of *Irx1* and *Scx* in the connective tissue between the MCPS and the PP further supports our hypothesis that this tissue forms through formation of a fibrocartilaginous joint.

## DISCUSSION

Sesamoid bones are small auxiliary bones that are highly variant in positioning, size and number (Bizarro, 1921; Jerez et al., 2009; Pearson and Davin, 1921b; Ponssa et al., 2010; Sarin et al., 1999). To date, the mechanism that regulates the development of these bones and allows the variations among them is missing. Here, by comparing the development of three different sesamoid bones, namely patella, lateral fabella and third metacarpophalangeal sesamoid, we identify common molecular and cellular mechanisms underlying these processes. Interestingly, we also identify points of diversion in their developmental program, which could provide a mechanistic explanation to the variability within this family of bones.

Our finding that all three studied sesamoid bones originated in *Sox9*- and *Scx*-positive precursors is interesting, because this lineage also contributes to the formation of bone superstructures, protrusions that project from the long bone surface and serve as attachment sites for tendons (Blitz et al., 2013; Sugimoto et al., 2013). These observations led us to suggest that bone superstructures and sesamoid bones can be placed under one category. Nevertheless, whereas bone superstructures protrude from the long bone shaft, sesamoid bones are auxiliary to it. Our finding that both patella and III-MCPS are separated from their bone of origin by a joint formation process offers a strategy for transforming a bone superstructure into a sesamoid bone, namely by detachment. Another way to determine between the development of a bone superstructure or a sesamoid bone is by regulating the spatial distribution of *Sox9^+^/Scx^+^* precursors. If the condensation of the *Sox9^+^/Scx^+^* precursors is adjacent to the target bone, then a bone superstructure will develop, as in the case of the deltoid tuberosity (Blitz et al., 2013). Conversely, if these cells condense away from the target bone, a sesamoid bone will be formed, as in the case of the lateral fabella.

An example for such transitional plasticity between sesamoid bones and superstructures is given by olecranon development. In mammalians, the olecranon forms as a bone superstructure at the proximal tip of the ulna, whereas in some anurans (frogs) it develops as a detached patella-like sesamoid bone (Ponssa et al., 2010), accordingly dubbed patella ulnaris. Presumably, formation of the olecranon as a sesamoid bone in earlier vertebrates preceded its formation as a fused superstructure in mammalians. Interestingly, studies in several mutated mouse lines have also demonstrated this developmental plasticity. Compound *Hox11^aadd^* mutant embryos were shown to exhibit a detached olecranon, similar to the patella ulnaris of anurans (Koyama et al., 2010). Another example was given by homozygous *Pbx1* mutant embryos, which developed a detached, sesamoid-like, deltoid tuberosity (DT) (Selleri et al., 2001).

The detachment mechanism has already been hypothesized over a century ago by Pearson and Davin (Pearson and Davin, 1921a; Pearson and Davin, 1921b), who suggested that sesamoid bones originate from existing bone eminences that detached from long bone shafts. F. G. Parsons proposed a contrary hypothesis suggesting that sesamoid bones develop independently yet could, in some cases, attach to neighboring long bones and become a bony eminence (Parsons, 1904; Parsons, 1908). Our data reconcile between these two opposing models by placing both bone eminences and sesamoid bones under one category of cartilage elements that develop from secondary *Sox9*^+^*/Scx*^+^ chondroprogenitors. Interestingly, our findings also suggest that both genetic and mechanical regulation are involved in facilitating developmental diversity, as indicated by both global and specific genetic regulation of the TGFβ/BMP signaling pathways and by the requirement of mechanical stimuli for patellofemoral joint formation. Our results show that specification of the precursors of all three sesamoids was regulated by TGFβ signaling and that BMP2 signaling regulated their growth, whereas BMP4 signaling was necessary for differentiation of patella precursors only. Moreover, whereas the lateral fabella formed as a separate entity, patella was separated by mechanical load-dependent formation of a synovial joint, and the third metacarpophalangeal sesamoid was separated by mechanical load-independent formation of a fibrocartilaginous joint. Altogether, these variations in the developmental programs of the different sesamoid bones may provide an initial explanation for the diversity in anatomy and function of this unique group of bones.

To conclude, our results provide further insight into the origin, formation and development of sesamoid bones. We suggest that these bony elements originate from pools of *Sox9*^+^/*Scx*^+^ chondroprogenitors whose regulation is independent from that of the long bone substructure and their induction is mechanical load-independent. We further show that some sesamoid bones form initially in juxtaposition to the long bone anlagen and are later separated by different programs of joint development. In contrast, we show that other sesamoids, such as the lateral fabella, can form independently of the long bones. Finally, we demonstrate that a combination of genetic and mechanical regulatory programs are applied to achieve anatomical and ultimately functional variation during sesamoid bones development.

## MATERIALS AND METHODS

### Animals

All experiments involving mice were approved by the Institutional Animal Care and Use Committee (IACUC) of the Weizmann Institute. For all timed pregnancies, plug date was defined as E0.5. For harvesting of embryos, timed-pregnant females were euthanized by cervical dislocation. The gravid uterus was dissected out and suspended in a bath of cold phosphate-buffered saline (PBS) and the embryos were harvested after removal of the placenta. Tail genomic DNA was used for genotyping by PCR.

Mice heterozygous for the mutation muscular dysgenesis (*mdg*) (Pai, 1965) were kindly provided by G. Kern (Innsbruck Medical University, Innsbruck, Austria). *Scx*-*Cre* and *ScxGFP* transgenic mice were obtained from R. Schweitzer (Shriners Hospital for Children Research Division, Portland, OR, USA) and R. L. Johnson (The University of Texas MD Anderson Cancer Center, Houston, TX, USA). C57/Bl6 mice (Jackson) were used for micro-CT analyses.

The generation of *Prx1*-*Cre* (Logan et al., 2002), floxed-*Tgfbr2* (Chytil et al., 2002), floxed-*Bmp4* (Liu et al., 2004; Selever et al., 2004), floxed-*Bmp2* (Ma and Martin, 2005), *ScxGFP* (Pryce et al., 2007) and *Rosa26-tdTomato* (Madisen et al., 2010) mice has been described previously. To create *mdg* mutant mice, animals heterozygous for the mutation were crossed; heterozygous embryos were used as a control. To create *Prx1*-*Tgfbr2, Prx1*-*Bmp4* or *Prx1*-*Bmp2* mutant mice, floxed-*Tgfbr2*, floxed-*Bmp4* or floxed-*Bmp2* mice were mated with *Prx1*-*Cre*-*Tgfbr2*, *Prx1*-*Cre*-*Bmp4* or *Prx1*-*Cre*-*Bmp2* mice, respectively. As a control, *Prx1*-*Cre*-negative embryos were used. For genetic lineage tracing analysis, *Scx*-*Cre* mice were crossed with homozygous *Rosa26-tdTomato* mice.

### Ex vivo micro-CT scanning

Before scanning, limbs of postnatal day (P) 14 C57/Bl6 mice (Jackson) were dissected, fixated overnight in 4% PFA/PBS and dehydrated to 100% ethanol. Samples were then scanned ex vivo in 100% ethanol by MicroXCT-400 (Xradia) at 30 kV and 4.5 W or 40 kV and 8 W.

### Paraffin sections

For preparation of paraffin sections, embryos were harvested at various ages, dissected and fixed in 4% paraformaldehyde (PFA)/PBS at 4°C overnight. After fixation, tissues were dehydrated to 100% ethanol and embedded in paraffin. The embedded tissues were cut to generate 7 μm-thick sections and mounted onto slides.

### OCT-embedded sections

For preparation of OCT-embedded sections, embryos were harvested at various ages, dissected and fixed in 1% paraformaldehyde (PFA)/PBS at 4°C overnight. Fixed embryos were then dehydrated gradually, first in 15% sucrose for 4-6 hours at room temperature and then in 30% sucrose overnight at 4°C. Next, samples were dissected and soaked in 15% sucrose/50% OCT for 30-60 minutes and then embedded in OCT. Frozen samples were immediately sectioned at a thickness of 10 μm and mounted onto slides.

### Histological analysis and fluorescent in situ hybridization

Safranin-O/Fast green staining was performed following standard protocols. Double fluorescent in situ hybridizations (dbFISH) on paraffin sections were performed using digoxigenin (DIG)-and fluorescein (FITC)-labeled probes (Shwartz and Zelzer, 2014). All probes are available upon request. Detection of probes was done by anti-DIG-POD (Roche, 11207733910, 1:300) and anti-FITC-POD (Roche, 11207733910, 1:300), followed by Cy3- or Cy2-tyramide labeled fluorescent dyes, according to the instructions of the TSA Plus Fluorescent Systems Kit (Perkin Elmer).

### Immunofluorescence staining

For immunofluorescence staining for SOX9 and collagen type II alpha 1 (COL2A1), 7 μm-thick paraffin sections of embryo limbs were deparaffinized and rehydrated to water. Antigen was then retrieved in 10 mM citrate buffer (pH 6.0), boiled and cooked for 10 minutes in a microwave oven. In order to block non-specific binding of immunoglobulin, sections were incubated with 7% goat serum, 1% BSA dissolved in PBST. Following blockage, sections were incubated overnight at 4°C with primary anti-SOX9 antibody (1:200; AB5535; Millipore). Then, sections were washed in PBST (PBS + 0.1% Triton X-100 + 0.01% sodium azide) and incubated with Cy3-conjugated secondary fluorescent antibodies (1:100; Jackson Laboratories). After staining for SOX9, slides were washed in PBST and fixed in 4% PFA at room temperature for 10 minutes. Then, slides were incubated with proteinase K (Sigma, P9290), washed and post-fixed again in 4% PFA. Next, sections were washed and incubated overnight at 4°C with primary anti-COL2A1 antibody (1:50; DSHB, University of Iowa). The next day, sections were washed in PBST and incubated with Cy2-conjugated secondary fluorescent antibodies (1:200; Jackson Laboratories). Occasionally, slides were counter-stained using DAPI. Slides were mounted with Immuno-mount aqueous-based mounting medium (Thermo).

For immunofluorescence staining for SOX9 and *ScxGFP*, 10 μm-thick cryostat sections of embryo limbs endogenously labeled for *ScxGFP* were used. SOX9 immunofluorescence staining was performed as described above, using primary SOX9 antibody and secondary Cy3-fluorescent antibodies.

## ACKNOWLEDGMENTS

We thank N. Konstantin for expert editorial assistance, S. Krief for expert technical support, and all other members of the Zelzer laboratory for their advice and suggestions. We thank R. Schweitzer for providing the *ScxGFP* mice. We thank C. Vega from the Weizmann Institute Department of Design, Photography and Printing for designing the model. This work was supported by grants from the United States-Israel Binational Science Foundation (BSF) [#2011122], the European Research Council (ERC) [#310098], and the Minerva Foundation [#711428].

## AUTHOR CONTRIBUTIONS

S.E. designed and carried out the experiments, analyzed the data and wrote the manuscript. S.R performed micro-CT experiments. S.K. jointly carried out experiments. L.L. performed Tgfb/Bmp experiments. E.Z. jointly designed the study, supervised the project, analyzed and interpreted the data, and wrote the manuscript.

## REFERENCES

Abdala, V., Vera, M.C. and Ponssa, M.L. (2017). On the presence of the patella in frogs. The Anatomical Record.

Askary, A., Mork, L., Paul, S., He, X., Izuhara, A. K., Gopalakrishnan, S., Ichida, J.K., McMahon, A.P., Dabizljevic, S., Dale, R., et al. (2015). Iroquois Proteins Promote Skeletal Joint Formation by Maintaining Chondrocytes in an Immature State. Developmental Cell 35, 358–365.

Bizarro, A.H. (1921). On Sesamoid and Supernumerary Bones of the Limbs. J Anat 55, 256–268.

Blitz, E., Sharir, A., Akiyama, H. and Zelzer, E. (2013). Tendon-bone attachment unit is formed modularly by a distinct pool of Scx - and Sox9 -positive progenitors. Development 140, 2680–2690.

Carter, D. (1998). Epigenetic mechanical factors in the evolution of long bone epiphyses. Zoological Journal of the Linnean Society 123, 163–178.

Chytil, A., Magnuson, M.A., Wright, C.V. and Moses, H.L. (2002). Conditional inactivation of the TGF□β type II receptor using Cre: Lox. Genesis 32, 73–75.

Corina Vera, M., Laura Ponssa, M. and Abdala, V. (2015). Further Data on Sesamoid Identity from Two Anuran Species: Morphology and Development of Anuran Skeletal Elements. The Anatomical Record 298, 1376–1394.

Doherty, A.H., Lowder, E.M., Jacquet, R.D. and Landis, W.J. (2010). Murine Metapodophalangeal Sesamoid Bones: Morphology and Potential Means of Mineralization Underlying Function. The Anatomical Record: Advances in Integrative Anatomy and Evolutionary Biology 293, 775–785.

Eyal, S., Blitz, E., Shwartz, Y., Akiyama, H., Schweitzer, R. and Zelzer, E. (2015). On the development of the patella. Development 142, 1831–1839.

Hall, B.K. (2005). Bones and cartilage: developmental and evolutionary skeletal biology. Academic Press.

Hauser, N.H., Hoechel, S., Toranelli, M., Klaws, J. and Müller-Gerbl, M. (2015). Functional and Structural Details about the Fabella: What the Important Stabilizer Looks Like in the Central European Population. BioMed Research International 2015, 1–8.

Jerez, A., Mangione, S. and Abdala, V. (2009). Occurrence and distribution of sesamoid bones in squamates: a comparative approach. Acta Zoologica.

Jin, Z.W., Shibata, S., Abe, H., Jin, Y., Li, X.W. and Murakami, G. (2017). A new insight into the fabella at knee: the foetal development and evolution. Folia Morphologica 76, 87–93.

Koyama, E., Yasuda, T., Minugh-Purvis, N., Kinumatsu, T., Yallowitz, A.R., Wellik, D.M. and Pacifici, M. (2010). Hox11 genes establish synovial joint organization and phylogenetic characteristics in developing mouse zeugopod skeletal elements. Development 137, 3795–3800.

Lennox, I.A., Cobb, A.G., Knowles, J. and Bentley, G. (1994). Knee function after patellectomy. A 12-to 48-year follow-up. Bone & Joint Journal 76, 485–487.

Liu, W., Selever, J., Wang, D., Lu, M.-F., Moses, K.A., Schwartz, R.J. and Martin, J.F. (2004). Bmp4 signaling is required for outflow-tract septation and branchial-arch artery remodeling. Proceedings of the National Academy of Sciences 101, 4489–4494.

Logan, M., Martin, J.F., Nagy, A., Lobe, C., Olson, E.N. and Tabin, C.J. (2002). Expression of Cre Recombinase in the developing mouse limb bud driven by a Prxl enhancer. Genesis 33, 77–80.

Ma, L. and Martin, J.F. (2005). Generation of a Bmp2 conditional null allele. genesis 42, 203–206.

Madisen, L., Zwingman, T.A., Sunkin, S.M., Oh, S.W., Zariwala, H.A., Gu, H., Ng, L.L., Palmiter, R.D., Hawrylycz, M.J., Jones, A.R., et al. (2010). A robust and high-throughput Cre reporting and characterization system for the whole mouse brain. Nature Neuroscience 13, 133–140.

Maisano, J.A. (2002). The potential utility of postnatal skeletal developmental patterns in squamate phylogenetics. Zoological Journal of the Linnean Society 136, 277–313.

McDonald, L.A., Gerrelli, D., Fok, Y., Hurst, L.D. and Tickle, C. (2010). Comparison of Iroquois gene expression in limbs/fins of vertebrate embryos: Iroquois gene expression. Journal of Anatomy 216, 683–691.

Mottershead, S. (1988). Sesamoid bones and cartilages: An enquiry into their function. Clinical Anatomy 1, 59–62.

Pai, A.C. (1965). Developmental genetics of a lethal mutation, muscular dysgenesis (MDG), in the mouse: I. Genetic analysis and gross morphology. Developmental biology 11, 82–92.

Parsons, F.G. (1904). Observations on traction epiphyses. Journal of anatomy and physiology 38, 248.

Parsons, F.G. (1908). Further remarks on traction epiphyses. Journal of anatomy and physiology 42, 388.

Pearson, K. and Davin, A.G. (1921a). On the Sesamoids of the Knee-Joint. Biometrika 13, 133.

Pearson, K. and Davin, A.G. (1921b). On the Sesamoids of the Knee-Joint: Part II. Evolution of the Sesamoids. Biometrika 13, 350.

Phukubye, P. and Oyedele, O. (2011). The incidence and structure of the fabella in a South African cadaver sample. Clinical Anatomy 24, 84–90.

Pignatti, E., Zeller, R. and Zuniga, A. (2014). To BMP or not to BMP during vertebrate limb bud development. Seminars in Cell & Developmental Biology 32, 119–127.

Ponssa, M.L., Goldberg, J. and Abdala, V. (2010). Sesamoids in Anurans: New Data, Old Issues. The Anatomical Record: Advances in Integrative Anatomy and Evolutionary Biology 293, 1646–1668.

Pryce, B.A., Brent, A.E., Murchison, N.D., Tabin, C.J. and Schweitzer, R. (2007). Generation of transgenic tendon reporters, ScxGFP and ScxAP, using regulatory elements of the scleraxis gene. Developmental Dynamics 236, 1677–1682.

Samuels, M.E., Regnault, S. and Hutchinson, J.R. (2017). Evolution of the patellar sesamoid bone in mammals. PeerJ 5, e3103.

Sarin, V.K., Erickson, G.M., Giori, N.J., Bergman, A.G. and Carter, D.R. (1999). Coincident development of sesamoid bones and clues to their evolution. The Anatomical Record 257, 174–180.

Schindler, O.S. and Scott, W.N. (2011). Basic kinematics and biomechanics of the patello-femoral joint Part 1□: The native patella. 77, 11.

Selever, J., Liu, W., Lu, M.-F., Behringer, R.R. and Martin, J.F. (2004). Bmp4 in limb bud mesoderm regulates digit pattern by controlling AER development. Developmental Biology 276, 268–279.

Selleri, L., Depew, M.J., Jacobs, Y., Chanda, S.K., Tsang, K.Y., Cheah, K.S., Rubenstein, J. L., O’Gorman, S. and Cleary, M.L. (2001). Requirement for Pbx1 in skeletal patterning and programming chondrocyte proliferation and differentiation. Development 128, 3543–3557.

Shukunami, C., Takimoto, A., Oro, M. and Hiraki, Y. (2006). Scleraxis positively regulates the expression of tenomodulin, a differentiation marker of tenocytes. Developmental biology 298, 234–247.

Shwartz, Y. and Zelzer, E. (2014). Nonradioactive in situ hybridization on skeletal tissue sections. In Skeletal Development and Repair, pp. 203–215. Springer.

Sugimoto, Y., Takimoto, A., Akiyama, H., Kist, R., Scherer, G., Nakamura, T., Hiraki, Y. and Shukunami, C. (2013). Scx+/Sox9+ progenitors contribute to the establishment of the junction between cartilage and tendon/ligament. Development 140, 2280–2288.

Sutton, F., Thompson, C., Lipke, J. and Kettelkamp, D. (1976). The effect of patellectomy on knee function: The Journal of Bone & Joint Surgery 58, 537–540.

Vickaryous, M.K. and Olson, W.M. (2007). Sesamoids and ossicles in the appendicular skeleton. Fins into limbs: evolution, development, and transformation 323–341.

Wirtschafter, Z.T. and Tsujimura, J.K. (1961). The sesamoid bones in the C3H mouse. The Anatomical Record 139, 399–408.

